# Adult-specific Reelin expression alters striatal neuronal organization. Implications for neuropsychiatric disorders

**DOI:** 10.1101/2023.01.21.525025

**Authors:** Mònica Pardo, Sara Gregorio, Enrica Montalban, Lluís Pujadas, Alba Elias-Tersa, Núria Masachs, Alba Vílchez-Acosta, Annabelle Parent, Carme Auladell, Jean-Antoine Girault, Miquel Vila, Angus C Nairn, Yasmina Manso, Eduardo Soriano

## Abstract

In addition to neuronal migration, brain development and adult plasticity, the extracellular matrix protein Reelin has been extensively implicated in human psychiatric disorders such as schizophrenia, bipolar disorder and autistic spectrum disorder. Moreover, heterozygous *reeler* mice exhibit features reminiscent of these disorders, while overexpression of Reelin protects against its manifestation. However, how Reelin influences the structure and circuits of the striatal complex, a key region for the above-mentioned disorders, is far from being understood, especially when altered Reelin expression levels are found at adult stages. In the present study, we took advantage of complementary conditional gain- and loss-of-function mouse models to investigate how Reelin levels may modify adult brain’s striatal structure and neuronal composition. Using immunohistochemical techniques, we determined that Reelin does not seem to influence the striatal patch and matrix organization (studied by μ-opioid receptor immunohistochemistry) nor the density of medium spiny neurons (MSNs, studied with DARPP-32). We show that overexpression of Reelin leads to increased numbers of striatal Parvalbumin- and Cholinergic-interneurons, and to a slight increase in the tyrosine hydroxylase-positive projections. We conclude that increased Reelin levels might modulate the numbers of striatal interneurons and the density of the nigrostriatal dopaminergic projections, suggesting that these changes may be involved in the protection of Reelin against neuropsychiatric disorders.

## 1 Introduction

Reelin is an extracellular matrix protein important for neuronal migration and layer formation during neocortical development (D’Arcangelo et al., 1995; Alcántara et al., 1998; Rice and Curran, 2001; Soriano and Del Río, 2005; Cooper, 2008; Hirota and Nakajima, 2017; Vílchez-Acosta et al., 2022). Besides its role during development, the Reelin pathway is also active in the adult brain, controlling glutamatergic neurotransmission, dendritic spine formation, synaptic plasticity and adult neurogenesis (Chen et al., 2005; Herz and Chen, 2006; Qiu et al., 2006b; Groc et al., 2007; Niu et al., 2008; Pujadas et al., 2010; Teixeira et al., 2012; Bosch et al., 2016). Reelin binds to Apolipoprotein E Receptor 2 (ApoER2) and Very-Low-Density Lipoprotein Receptor (VLDLR), leading to the phosphorylation and activation of the intracellular adaptor protein Disabled 1 (Dab1), which triggers a complex signalling cascade involving members of the Src kinase family, the PI3K, Erk1/2 and GSK3 kinases, and Cullin-5-dependent degradation, amongst others (Howell et al., 1997, 1999; D’Arcangelo et al., 1999; Hiesberger et al., 1999; Beffert et al., 2002; Arnaud et al., 2003; Benhayon et al., 2003; Ballif et al., 2004; Strasser et al., 2004; González-Billault et al., 2005; Simó et al., 2007, 2010; Yasui et al., 2010; Molnár et al., 2019).

Genetic studies have associated the Reelin gene (RELN) with a number of psychiatric diseases, including schizophrenia, bipolar disorder and autistic spectrum disorder (Impagnatiello et al., 1998; Fatemi et al., 2001, 2005; Persico et al., 2001; Grayson et al., 2005; Ovadia and Shifman, 2011; Wang et al., 2014; Baek et al., 2015; Lammert and Howell, 2016). This link is also supported by studies showing that Reelin levels are reduced in patients with schizophrenia and bipolar disorder (Fatemi et al., 2000; Torrey et al., 2005; Ruzicka et al., 2007), and can be altered by psychotropic medication (Fatemi et al., 2009). In fact, Reelin haploinsufficiency models, based on the suppression or reduction of Reelin expression (or its downstream pathway), manifest features related to neuropsychiatric disorders, such as cognitive impairments, psychosis vulnerability and learning deficits that frequently coexist with evident alterations in hippocampal plasticity(Tueting et al., 1999; Krueger et al., 2006; Marrone et al., 2006; Qiu et al., 2006a; Ammassari-Teule et al., 2009; Folsom and Fatemi, 2013). Conversely, overexpression of Reelin protects against psychiatric disease-related phenotypes in mice, since it reduces cocaine sensitization, disruption of prepulse inhibition (PPI) and the time spent floating in the forced swim test (Teixeira et al., 2011). Furthermore, Reelin also regulates adult neurogenesis and synaptogenesis (Kim et al., 2002; Pujadas et al., 2010; Teixeira et al., 2012; Bosch et al., 2016), whose disruption is considered to be involved in the pathogenesis of psychiatric disorders (Kempermann, 2008; Zhao et al., 2008).

The striatum plays a critical function in motor control and regulation of motivated behaviours (Bolam et al., 2000). Its neuronal population is composed by a 5-10% of interneurons and the rest (90-95%) are GABAergic medium spiny neurons (MSNs). The latter can be classified into striatonigral or striatopallidal subtypes based on their axonal projections to the internal Globus pallidus (iGP) and Substantia Nigra (SN) or to the external Globus Pallidus (eGP), respectively. They can be distinguished by the expression of the Dopamine D1 receptor (striatonigral MSNs) or the Dopamine D2 receptor (striatopallidal MSNs) (Bolam, 1984; Schiffmann et al., 1991; Gerfen, 1992; Smith et al., 1998). Although the striatum exhibits a relatively uniform appearance, it presents a complex organization based in two different compartments: the patches or striosomes (stained by μ -opioid receptor MOR) and the matrix, which surrounds the patches (Olson et al., 1972; Graybiel and Ragsdale, 1978; Herkenham and Pert, 1981). A proper cellular and compartmental organization is essential for a correct striatal function (Crittenden and Graybiel, 2011).

Besides the involvement of the striatum (including the Nucleus accumbens) and its circuitry in psychiatric disorders such as major depression, schizophrenia and obsessive-compulsive disorder (OCD), few studies addressing how Reelin influences striatal structure and circuits are available (de Guglielmo et al., 2022). Most of these studies use heterozygous reeler mice as a model, which have reduced Reelin expression also during development. Here we investigate how altering Reelin levels, specifically at late postnatal and adult stages, may lead to cellular and compartmental changes in the striatum that could be related to neuropsychiatric disorders. We used gain- and loss-of-function conditional mouse models to investigate how Reelin levels may modify striatal structure and neuronal composition. Our results suggest that whereas Reelin does not seem to influence the patch-matrix striatal organization and the numbers of MSNs, overexpression of Reelin leads to increased numbers of striatal interneurons and to a slight increase in the dopaminergic projections.

## 2 Materials and methods

### 2.1. Animals

The TgRln is a conditionally regulated transgenic line that overexpresses Reelin by a transactivator (tTA) under the control of the calcium–calmodulin-dependent kinase II α promoter (pCaMKIIα)(Pujadas et al., 2010). Reelin transgenic littermates, which have an inactive form of the Reelin gene insertion without the transactivator tTA, were used as controls. For the generation of the Reelin conditional knockout mouse line, homozygous floxed Reelin (fR/fR) mice, with the exon 1 of the Reln gene flanked by loxP sites, were crossed with a heterozygous UbiCreERT2 line (B6.Cg-Tg(UBC-cre/ESR1)1Ejb/J, stock #008085, The Jackson Laboratory), both on a C57BL/6J background (Vílchez-Acosta et al., 2022). The UbiCreERT2 line displays a ubiquitous expression of the Cre recombinase fused to a modified estrogen receptor ligand-binding domain that retains the Cre at the cytoplasm. Administration of an estrogen receptor antagonist (tamoxifen) induces the nuclear translocation of Cre recombinase and the ubiquitous scission of the floxed gene sequence (Reln) in all tissues. The resultant offspring (Cre fR/fR) was used for the experiments, and fR/fR littermates were used as controls. In both transgenic lines, 4-5 months old female and male mice were used for the experiments.

Male, 8–10-week old, Drd2-EGFP (n=20 Swiss-Webster and 6 C57BL/6N background, founder S118), Drd1a-EGFP (n=4 Swiss-Webster and n=4 C57BL/6N background, founder X60) hemizygous mice were also used in this study. BAC Drd2- and Drd1a-EGFP mice, that express the reporter protein enhanced green fluorescent protein under the control of the D2 and D1 receptor promoters, were generated by GENSAT (Gene Expression Nervous System Atlas) at the Rockefeller University (New York, NY)(Gong et al., 2007).

Mice were bred, studied and processed at the animal research facility of the Faculty of Pharmacy of the University of Barcelona and at the animal research facility of the Rockefeller University. Animals were provided with food and water ad libitum and maintained in a temperature-controlled environment in a 12/12 h light-dark cycle. All the experiments involving animals were performed in accordance with the European Community Council directive 2010/63/EU, the National Institute of Health guidelines for the care and use of laboratory animals, and the Rockefeller University’s Institutional Animal Care and Use Committee (protocol 14753-H). Experiments were also approved by the local ethical committees.

### 2.2. PCR Genotyping

DNA was extracted from tail biopsies by adding 100μl Sodium Hydroxide (50mM), and incubating at 100ºC during 15 minutes. Then, samples were kept on ice for 10 minutes and stored at -20ºC until use.

The PCR was performed with the GoTaq® Green Master Mix (Promega), and the primers used for genotyping were as follows. Cre fR/fR line: for homozygous floxed Reelin detection, FloxA (5’CGAGGTGCTCATTTCCCTGCACATTGC3’) and FloxB (5’ CACCGACCAAAGTGCTCCAATCTGTCG 3’) primers were used. Homozygous fR/fR mice present only one band of 613 bp whereas heterozygous mice present an additional band at 496 bp. To determine the presence of UbiCre, the primers UbiCre1(5’ GCG GTC TGG CAG TAA AAA CTA TC 3’) and UbiCre2 (5’ GTC AAA CAG CAT TGC TGT CAC TT 3’) which are specific for UbiCreERT2, and UbiCre3 (5’ CTA GGC CAC AGA ATT GAA AGA TCT 3’) and UbiCre4 (5’ GTA GGT GGA AAT TCT AGC ATC ATC C 3’) as internal positive control were used. Mice heterozygous for Cre (Cre fR/fR) had a double band at 324 and 100 bp while mice negative for Cre only amplified the 100 bp band. TgRln line: the primers RLTG-gen-F (5’-TTGTACCAGGTTCCGCTGGT-3’) and RLTG-gen-R (5’-GCA CAT ATC CAG GTT TCA GG-3’) were used to amplify both the endogenous Reelin gene (720bp) and the transgenic DNA (320 bp); the primers nTTA-C (5’-ACT AAG TCA TCG CGA TGG AG-3’) and nTTA-F (5’-CGA AAT CGT CTA GCG CGT C-3’), were used to detect the transactivator tTA transgene (Pujadas et al., 2010).

### 2.3. Tamoxifen administration

Inactivation of Reelin expression was induced at postnatal day (p)45-60 by daily intraperitoneal injections of tamoxifen dissolved in 10% alcohol-90% sunflower oil for 3 consecutive days (180mg/kg/day; Sigma-Aldrich).

### 2.4. Immunohistochemistry

For immunohistochemistry, 4-5 months old mice were perfused transcardially with 4% paraformaldehyde (PFA) in PB 0.1M. Brains were quickly removed, fixed overnight in PFA, and then transferred to 30% sucrose in PBS 0.1M and stored at 4 ºC (48h). Brains were frozen with methylbutane (Honeywell) at -42ºC and stored at -80ºC until use. Thirty-μm coronal sections were obtained with a freezing microtome (Leica SM2010R) and were kept in a cryoprotective solution at - 20ºC. Immunohistochemistry was performed on free-floating sections. The sections were inactivated for endogenous peroxidases with 3% H_2_O_2_ in 10% Methanol and PBS for 15 minutes. After 3 washes with PBS and 3 washes with PBS-0.2%Triton (PBS-T), sections were blocked for 2 h at room temperature (RT) with PBS-T containing 10% of normal horse serum (NHS) and 0.2% of gelatin. For Reelin immunostaining, anti-mouse unconjugated F(ab’)2 fragments (1:300, Jackson ImmunoResearch), were added in the blocking step. After 3 washes with PBS-T, tissue sections were incubated with a primary antibody with PBS-T containing 5% of NHS and 0.2% of gelatine, overnight at 4ºC.

The commercial primary antibodies used were: anti-Reelin (clone G10, MAB5364, Merck Millipore, 1:1,000), anti-Choline Acetyltransferase (ChAT AB144P, Merck Millipore,1:500,), anti-μ Opioid Receptor (MOR, 1:2000, rabbit, AB5511, Merck Millipore), anti-Parvalbumin (PV, 1:500, Rabbit, PV27, Swant), anti-Dopamine- and cAMP-regulated phosphoprotein, 32 kDa (Darpp32, 1:500, mouse, 611520, BD Transduction Laboratories), anti-Tyrosine Hydroxilase (TH, 1:1000, Rabbit, AB152, Merck Millipore). Sections were washed with PBS-T and then incubated for 2 h at RT with biotinylated secondary antibody (1:200, Vector Laboratories). After subsequent washes with PBS-T, the sections were incubated for 2h at RT with streptavidin-HRP (1:400, GE Healthcare UK). After washing, the staining was developed using 0.03% diaminobenzidine (DAB) and 0.01% H_2_O_2_, with 0.1% nickel ammonium sulphate added to the solution. Finally, sections were dehydrated and mounted with Eukitt mounting medium (Sigma-Aldrich).

For immunofluorescence staining a similar procedure was followed using AlexaFluor 488 secondary antibody (1:500, Invitrogen, ThermoFisher) (excluding peroxidase inactivation), counterstained with Bisbenzimide (1:500) for 30 minutes at RT, mounted with Mowiol and stored at -20ºC.

### 2.5. D1-/D2-cell specific mRNA extraction

Cell-type specific translated-mRNA purification (TRAP), was performed as previously described (Heiman et al., 2008) with a few modifications. Each sample consisted of a pool of 2-3 mice. BAC-TRAP transgenic mice (Drd2- and Drd1a-EGFP) were sacrificed by decapitation. The brain was quickly dissected out and placed in a cold buffer and was then transferred to an ice-cold mouse brain matrix to cut thick slices from which the Nucleus Accumbens (NAcc) and the Dorsal Striatum (DS) were punched out using ice-cold stainless-steel cannulas. Each sample was homogenized in 1 ml of lysis buffer (20 mM HEPES KOH [pH 7.4], 5 mM MgCl2, 150 mM KCl, 0.5mM dithiothreitol, 100 μg/ml CHX protease and RNAse inhibitors) with successively loose and tight glass-glass 2 ml Dounce homogenizers. Each homogenate was centrifuged at 2000 x g, at 4°C, for 10 min. The supernatant was separated from cell debris and supplemented with NP-40 (EDM Biosciences) to a final concentration of 1% and DHPC (Avanti Polar lipids) to a final concentration of 30 mM. After mixing and incubating on ice for 5 minutes, the lysate was centrifuged for 10 minutes at 20,000 x g to separate the supernatant from the insolubilized material. A mixture of streptavidin-coated magnetic beads was incubated with biotinylated protein L and then with GFP antibody that was added to the supernatant and incubated ON at 4°C with gentle end-over rotation. After incubation, beads were collected with a magnetic rack and washed 5 times with high-salt washing buffer (20 mM HEPES-KOH [pH 7.4], 5 mM MgCl2, 150 μl 1M, 350 mM KCl, 1% NP-40) and immediately placed in “RTL plus” buffer (Qiagen). The mRNA was purified using the RNase micro KIT (Qiagen). RNA integrity was checked with the Bionalyzer (agilent 2100 Bioanalyzer, Agilent RNA 6000 nano kit). Five ng of mRNA from each sample were used for retro-transcription, performed with the Reverse Transcriptase III (Life Technologies) following the manufacturer’s instructions.

### 2.6. Real-Time PCR

Quantitative real time PCR, was performed using SYBR Green PCR kit in 96-well plates according to the manufacturer’s instructions. Results are presented as normalized to the indicated house-keeping genes and the delta-threshold cycle (Ct) method was used to obtain a fold change. mRNA levels are presented relative to D2. The housekeeping gene for normalization was beta-myosin heavy chain gene (Myh7).

### 2.7. Immunohistochemical analysis

For Darpp-32 cell counting, sections were scanned using NanoZoomer 2.0-HT (Hamamatsu). We used FIJI software to crop the striatal profile from the image. Darpp-32 positive cells were counted with the cell nuclei assistant TMarker software.

The images of PV and ChAT interneurons were acquired with a Nikon E600 microscope attached to an Olympus DP72 camera, and images were reconstructed using MosaicJ from the Fiji software (Fiji is Just Image J - NIH). The intermediate striatum was subdivided into four sub-regions: Dorso-Medial (DM), Dorso-Lateral (DL), Ventro-Medial (VM) and Ventro-Lateral (VL) (see (Gernert et al., 2000; Ammassari-Teule et al., 2009)) taking slices from Bregma 1.34 mm to 0.02 mm, to identify possible changes in the neuronal distribution inside the different striatal regions. Cell density studies were performed with FIJI tools to measure the area and to count cells (cell counter).

To measure TH intensity, slides were scanned with SilverFast at 600ppm and SigmaPlot was used to measure the intensity of the different striatal areas. The results are expressed as % from control which was considered as 1 in each independent experiment to avoid deviations caused by differences in the DAB development procedure.

Synaptic bouton images were taken with 63X oil immersion objective and counted selecting randomly an 11×11 mm^2^ ROI using Fiji.

For each mouse transgenic line we analyzed 3-14 animals and for each animal and average of 6-8 images were analyzed.

### 2.8. Statistics

All statistical analyses were performed using Graphpad Prism 5.0 software (GraphPad Software, Inc). Data was analyzed with unpaired two-tailed Student’s T-tests and statistical significance was set at p-value <0.05. Unless otherwise stated, all values are presented as mean ± the standard error of the mean (SEM). The number of animals used in each experiment is detailed in the figure legend.

## 3 Results

### 3.1. Reelin is highly expressed in striatonigral MSNs

To determine the effects of Reelin levels in the mouse striatal organization, we first studied Reelin expression in a Reelin overexpressing and a knock out mouse line. Control mice from both lines exhibited numerous Reelin-positive cell bodies that were distributed throughout the striatum (Figure 1A, C), whereas the tamoxifen-inducible conditional knockout mouse line (Cre fR/fR) presented a drastic reduction of Reelin protein as detected by immunohistochemistry (Figure 1B) and by western blot (not shown). In contrast, Reelin overexpressing mice (TgRln) showed a dramatic increase of Reelin protein in the striatum (Figure 1D) which was apparent in both the cell bodies and in the neuropil (see also (Pujadas et al., 2010)).

**Figure 1.**
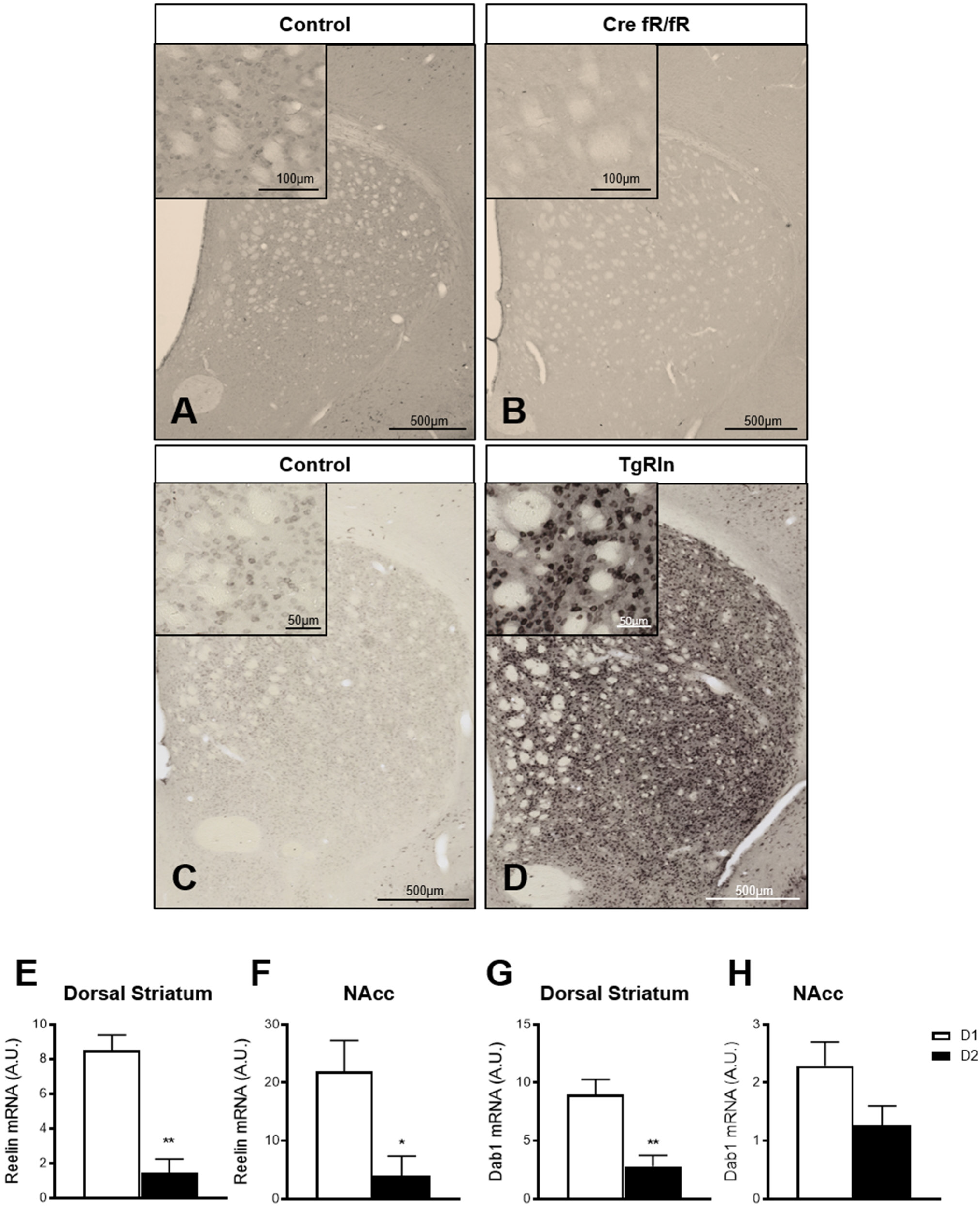
Reelin in the striatum is mainly expressed by D1 striatonigral MSNs. Immunohistochemistry against Reelin shows that Reelin protein is absent in the striatum of Cre fR/fR mice (**B**) compared to the controls (**A**) while it is clearly overexpressed in the striatum of TgRln mice (**D**) compared to controls (**C**). Quantification of Reelin mRNA levels in the Dorsal striatum (**E**) and NAcc (**F**) of D1/D2-TRAP mice (n=3-4). Quantification of Dab1 mRNA levels in the Dorsal striatum (**G**) and NAcc (**H**) of D1/D2-TRAP mice (n=4-7). NAcc, Nucleus Accumbens; D1, Dopamine 1 Receptor; D2, Dopamine 2 Receptor. Statistical analysis was performed using Student’s t-test; significant differences were established at *p<0.05, **p<0.01. Data represents mean±SEM.

Reelin has been described to co-localize with Calbindin D-28k-positive neurons (Sharaf et al., 2015), a well-known marker of striatal MSNs. Hence, we used the TRAP technology (Heiman et al., 2008) to determine a possible enrichment of Reelin mRNA in D1- or D2-receptor expressing MSNs in both DS and NAcc. BAC-TRAP-D1 and -D2 mice, were used to specifically immunoprecipitate mRNAs from D1 (striatonigral) or D2 (striatopallidal) neuronal populations from the DS and the NAcc. Reelin mRNA levels were compared to the housekeeping beta-myosin heavy chain gene. Results indicated that Reelin mRNA is enriched in D1-MSNs, in both the DS and the NAcc (Figure 1E, F). The expression of Dab1, a key downstream effector of the Reelin pathway, was also higher in D1 MSNs of the DS and NAcc (Figure 1G, H). These results suggest that the striatonigral D1 MSNs population is the main producer of striatal Reelin.

### 3.2. Striatal MSNs organization is independent of Reelin expression levels

To determine whether Reelin expression levels could modify DS MSN populations, we first immunostained sections with Darpp-32, a marker of MSNs, and quantified the density of striatal MSNs in the Cre fR/fR (Figure 2A, B) and TgRln (Figure 2C, D) mouse models. Results indicated that neither the absence nor the overexpression of Reelin altered the density of striatal Darpp-32 positive neurons in the striatum of Cre fR/fR (Figure 2A-B, E) or TgRln mice (Figure 2C-D,F).

**Figure 2.**
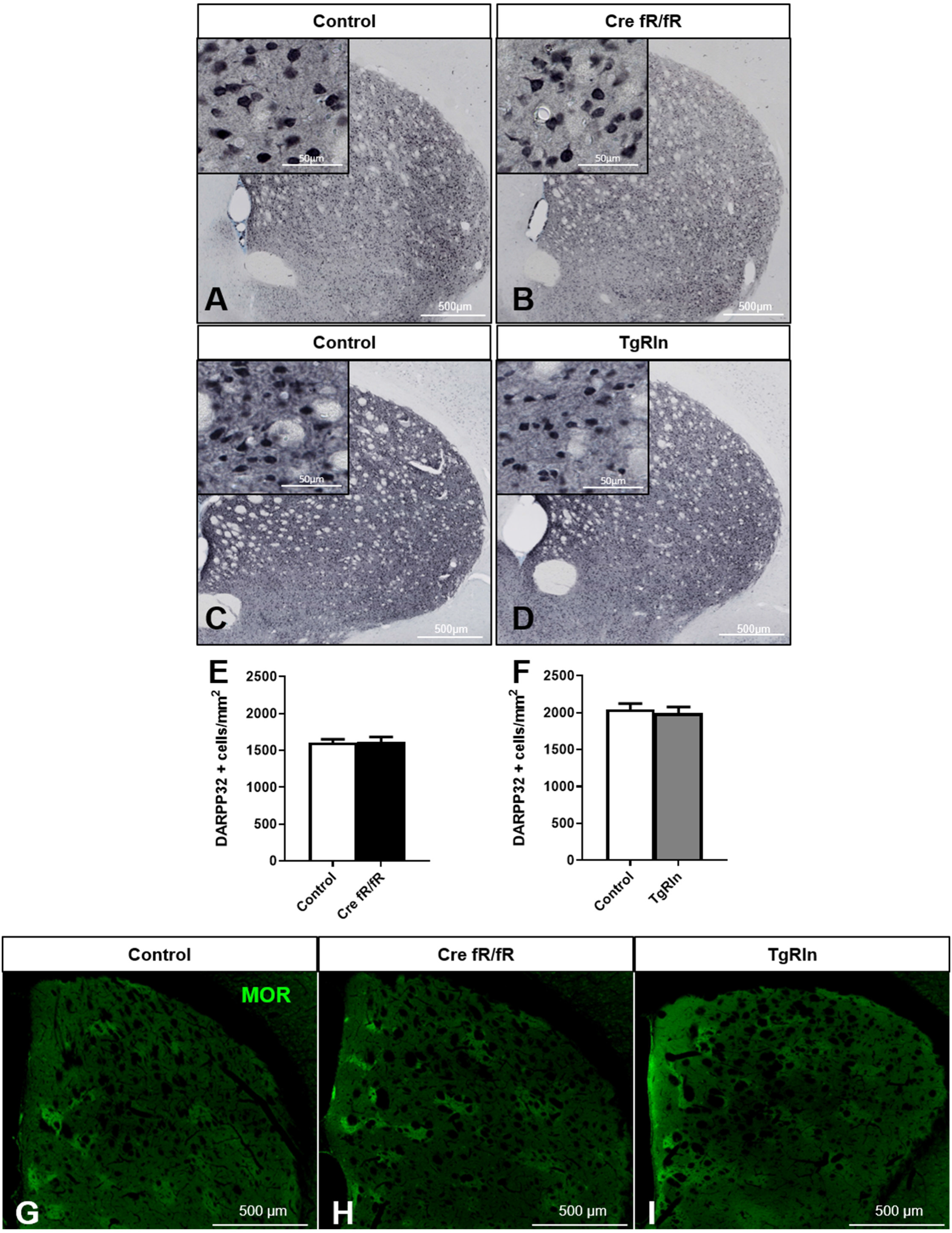
Striatal MSNs density and organization is not affected by Reelin levels. Representative images of Darpp-32 immunohistochemistry (striatal MSNs) in coronal sections of control and Cre fR/fR (**A-B**) and control and TgRln (**C-D**) striatum. Quantification of Darpp-32 cell density showed no alterations of striatal MSNs neither in Cre fR/fR (n=5-6) (**E**) nor in TgRln (n=6) **(F)** mice. Immunofluorescence for μ-Opioid receptor (MOR) in coronal sections of control (**G**), Cre fR/fR (**H**) and TgRln (**I**) striatum showing a similar organization of striatal patches in all the models. Statistical analysis was performed using Student’s t-test. Data is represented as mean±SEM.

Since Reelin controls neuronal migration, we next wanted to determine whether Reelin levels could affect the DS patch organization. Immunostaining of the striosomes with MOR showed striatal patches with a similar spatial distribution in all genotypes, suggesting that striatal MSNs density and organization are not affected by alterations of Reelin expression levels (Figure 2G-I).

### 3.3. Reelin overexpression alters striatal interneuron population

In addition to MSNs, the striatum also contains ChAT-positive and GABAergic interneurons, being the PV-expressing ones the best known. To assess the number and distribution of ChAT-positive interneurons in the different transgenic lines, we subdivided the DS in four Dorso-Ventral and Medio-Lateral regions (Figure 3A). Analysis of the density and distribution of ChAT-positive cells showed no differences in Cre fR/fR mice compared to controls (Figure 3A-F). In contrast, the density of ChAT-positive cells was increased in Reelin overexpressing mice compared to controls, reaching significance in 3 of the striatal sub-regions analyzed (Figure 3G-L)

**Figure 3.**
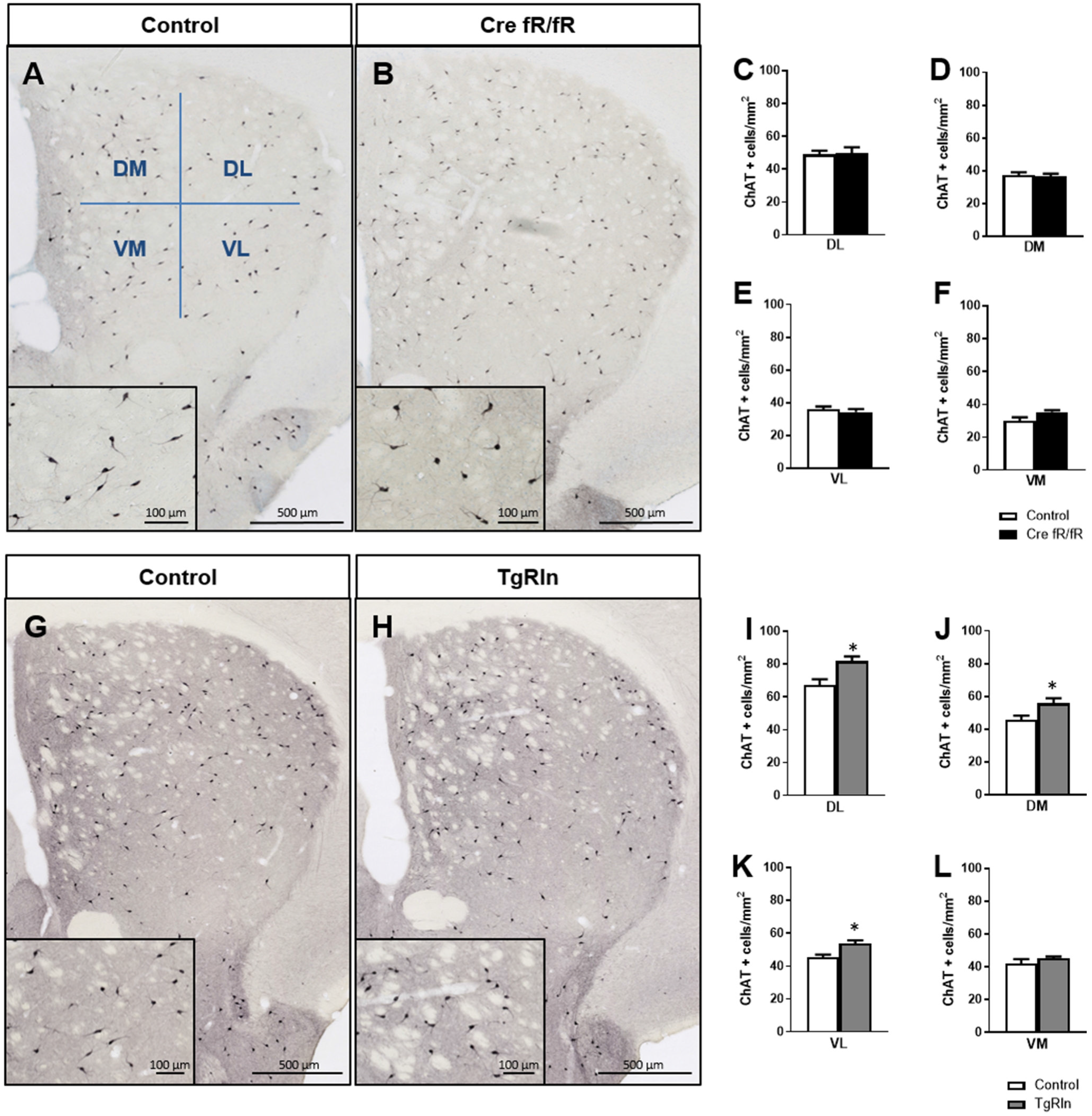
Reelin overexpression increased the density of striatal cholinergic interneurons. Immunohistochemistry of ChAT in striatal coronal sections of control and Cre fR/fR mice (**A-B**), with representative subdivision of the striatum in four regions (DM, Dorsal-Medial; DL, Dorsal-Lateral; VM, Ventral-Medial; VL, Ventral-Lateral). Quantification of ChAT density in the striatal subdivisions showed no differences between control and Cre fR/fR mice (n=4) (**C-F**). Representative images of ChAT immunohistochemistry in the striatum of control and TgRln mice, with higher magnification insets showing increased ChAT+ neuronal density in the TgRln mice (**G-H**). Quantification of ChAT cell density indicated a significant increase in the DL, VL and DM striatal regions of TgRln mice (n=4-6) (**I-L**). Statistical analyses were performed using Student’s t-test; *p<0.05. Data is represented as mean±SEM.

We also analyzed the density and distribution of PV striatal interneurons. In line with the ChAT-positive interneuron data, no changes in the density and distribution of PV-positive interneurons (Figure 4A-B) were observed in any of the DS regions of Cre fR/fR mice compared to controls (Figure 4C-F). However, analysis of PV-positive interneurons density in TgRln mice showed a statistically significant increase in the VL striatum (Figure 4G, H, K) but not in other striatal regions (Figure 4 G-J, L) as compared to controls. Altogether, our results indicated that Reelin overexpression increased the number of DS interneurons.

**Figure 4.**
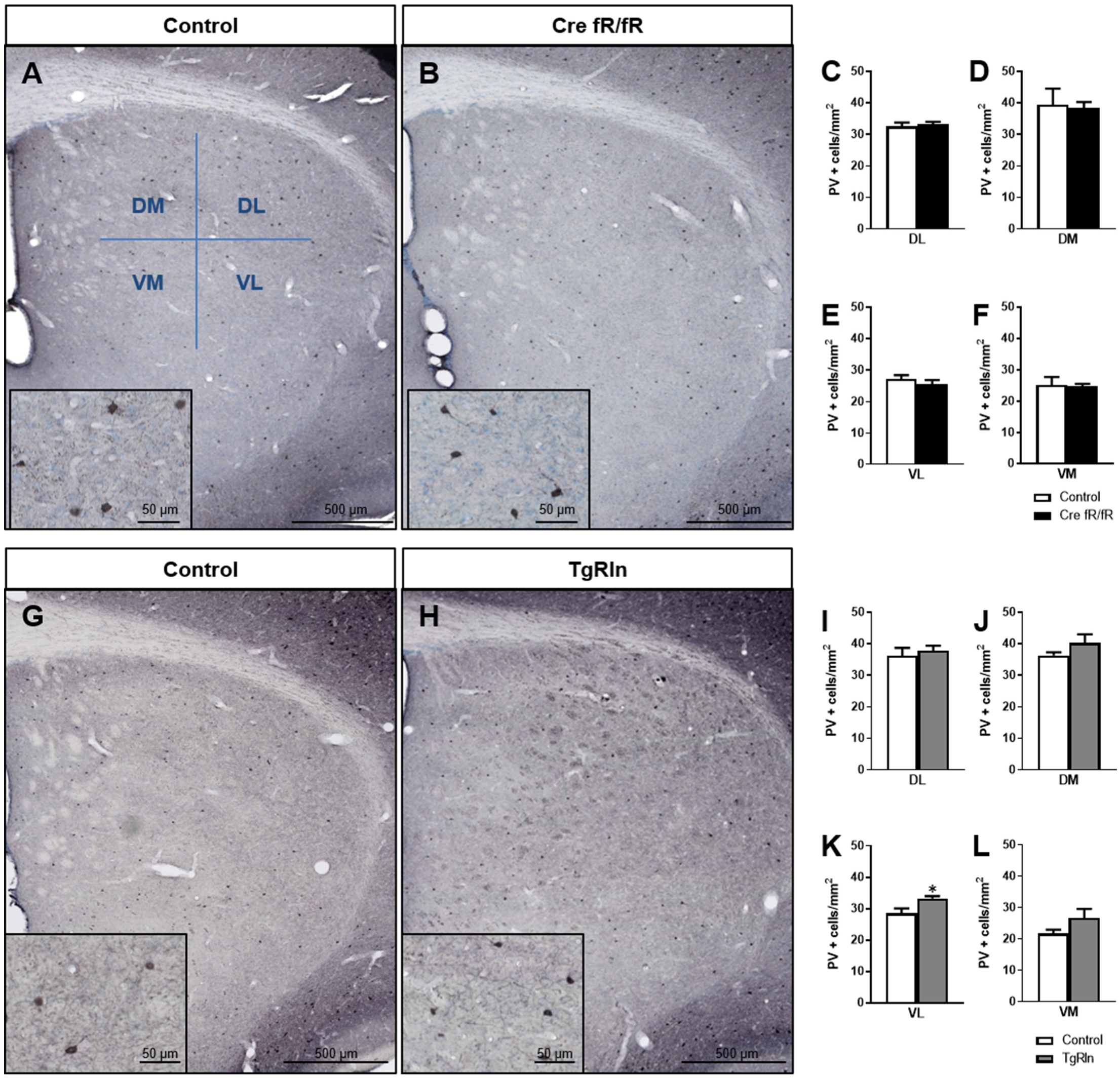
Increased levels of Reelin alter the density of Parvalbumin interneurons in the ventral-medial striatum. Immunohistochemistry for PV in coronal sections of control and Cre fR/fR striatum (**A-B**), subdividing the striatum in four regions (DM, Dorsal-Medial; DL, Dorsal-Lateral; VM, Ventral-Medial; VL, Ventral-Lateral). Quantification of the density of PV interneurons indicated no differences between the control and Cre fR/fR mice (n=4) (**C-F**). Representative images of PV immunostaining in the striatum of control and TgRln mice (**G-H**). Quantification of PV immunohistochemistry indicated an increase in the density of PV positive cells in the VL striatum of TgRln mice (n=4-5) (**K**) with no differences in the other striatal regions (**I-J, L**). Statistical analyses were performed using Student’s t-test; *p<0.05. Data is represented as mean±SEM.

### 3.4. Reelin levels control dopaminergic projections

Next, we analyzed whether the expression of Reelin could influence dopaminergic projections. Thus, we performed immunohistochemistry for TH to detect dopaminergic projections that reach the striatum from the Substantia Nigra (SN) and the Ventral Tegmental Area (VTA). We quantified TH intensity in the DS and the Ventral Striatum (VS), including the NAcc and the Olfactory Tubercle (OT). In the Cre fR/fR model, we observed no alterations in the dopaminergic intensity in none of the three striatal regions studied (Figure 5A-E) compared to controls. However, in the OT of Cre fR/fR mice, we observed a tendency towards a reduction in TH intensity compared to controls (Figure 5E). In contrast, in Reelin overexpressing mice, quantification of TH immunostaining (Figure 5F-G) showed a significant increase of TH intensity in both the NAcc and OT regions compared to controls (Figure 5H-J).

**Figure 5.**
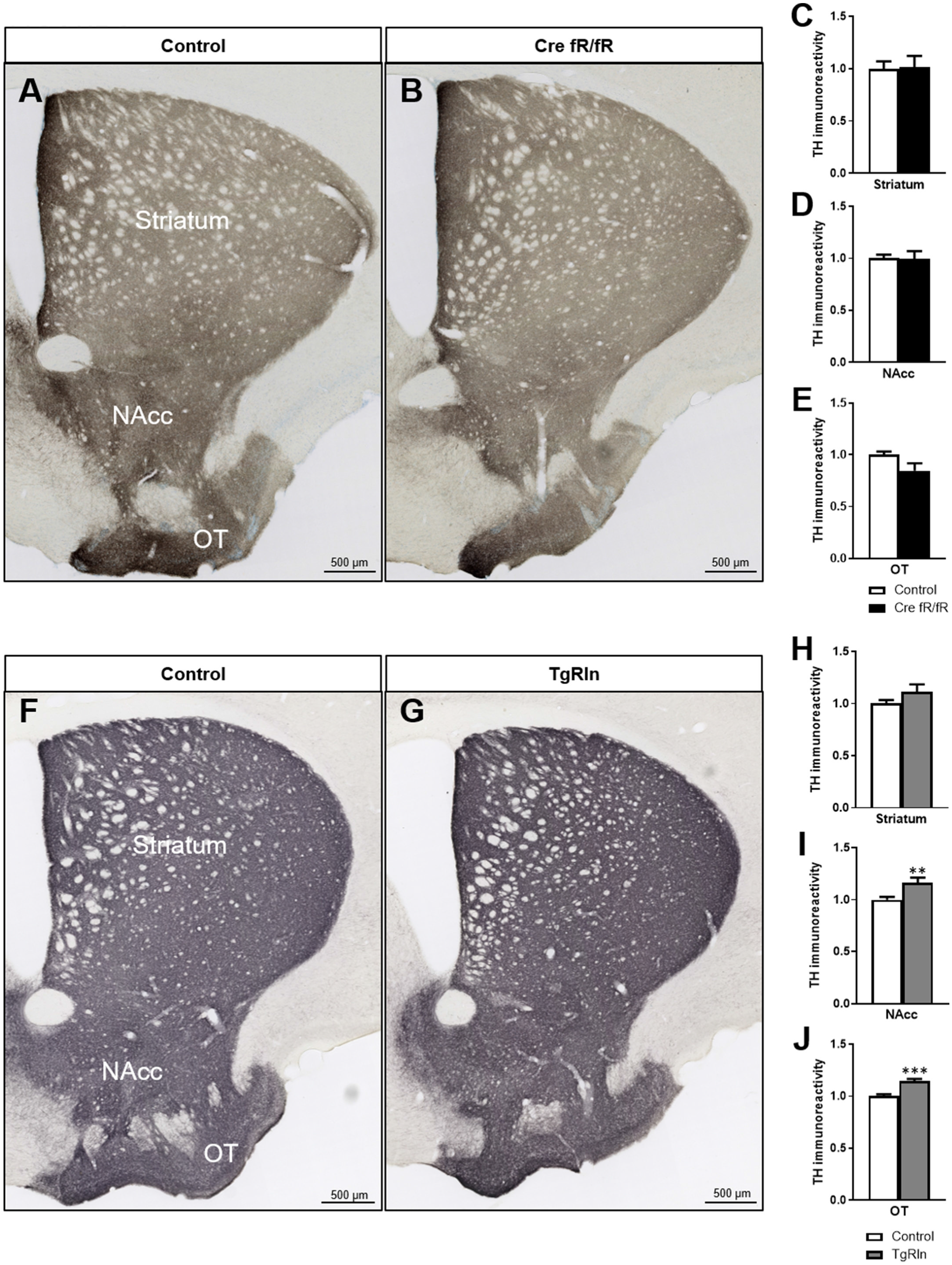
Increase of Reelin expression elevates dopaminergic projections in the Ventral Striatum. Immunohistochemistry for TH to stain dopaminergic projections in coronal sections of the DS, NAcc and OT of control and Cre fR/fR mice (**A-B**). TH intensity remains constant in the striatum (**C**), NAcc (**D**) and OT (**E**) of Cre fR/fR mice compared to controls (n=4) (**C-E**). Immunohistochemistry for TH in control and TgRln mice (**F-G**). Increased TH immunoreactivity was detected in the NAcc (**I**) and OT (**J**) but not in the DS (**H**) of TgRln mice compared to controls (n=8-14). NAcc, Nucleus Accumbens; OT, Olfactory Tubercle. Statistical analyses were performed using Student’s t-test; **p<0.01; ***p<0.001. Results represent the mean±SEM.

Finally, we also wanted to quantify synaptic boutons of striatal dopaminergic projections. Thus, we determined the density of synaptic boutons in the DS, NAcc and OT, dividing the DS into Dorsal and Ventral regions. In the Cre fR/fR mice, the density of synaptic boutons in all the regions was similar to that of control mice (DS dorsal: fR/fR 0.2838±0.017 vs. Cre fR/fR 0.3025±0.014; DS ventral: fR/fR 0.2720±0.016 vs. Cre fR/fR 0.2688±0.023; NAcc: fR/fR 0.2618±0.013 vs. Cre fR/fR 0.2530±0.034; OT: fR/fR 0.2428±0.013 vs. Cre fR/fR 0.2493±0.018; n= 4 mice/genotype, Mean±SD). In contrast, the density of dopaminergic synaptic boutons tended to increase in the TgRln mice compared to controls (Figure 6A-L), being statistically significant in the NAcc (Fig. 6C, G, K). These results suggest that higher Reelin levels might modulate dopaminergic fibres and synaptic boutons, mainly in the NAcc.

**Figure 6.**
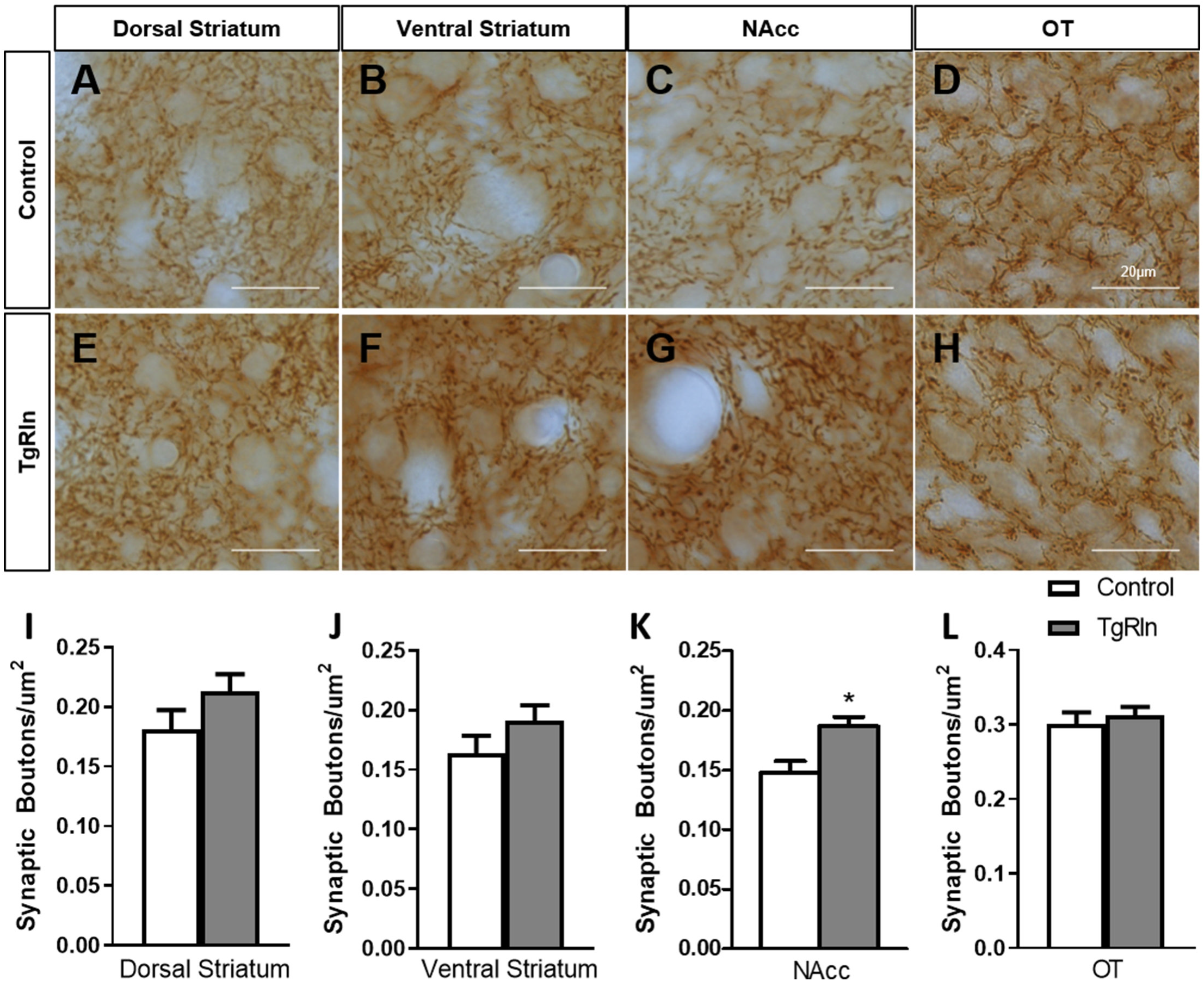
Increased number of dopaminergic synaptic boutons in the NAcc of TgRln mice. Immunohistochemistry for TH staining dopaminergic synaptic boutons in the dorsal (**A, E**) and ventral regions (**B, F**) of the DS, NAcc (**C, G**) and OT (**D, H**) of TgRln mice and its controls. Quantification of the density of dopaminergic boutons evidenced a higher density of synaptic boutons in the NAcc (**K**) of TgRln mice compared to its controls while no differences were observed in the rest of the analysed structures (**I-J, L**). DS, Dorsal striatum; NAcc, Nucleus Accumbens; OT, Olfactory Tubercle. Statistical analyses were performed using Student’s t-test; *p<0.05. Data is represented as mean±SEM.

## 4 Discussion

Variations in Reelin expression levels have been shown to be important for the development of neuropsychiatric disorders (Impagnatiello et al., 1998; Fatemi et al., 2000, 2001, 2005; Persico et al., 2001; Torrey et al., 2005; Grayson et al., 2005; Ruzicka et al., 2007; Ovadia and Shifman, 2011; Wang et al., 2014; Baek et al., 2015; Lammert and Howell, 2016); however, we still lack the precise understanding of the mechanistic insights of this correlation. Here we focused our attention on the striatum as a key region participating in the pathogenesis of psychiatric diseases (McCutcheon et al., 2021). We thus characterized specific striatal neuronal populations as well as the dopaminergic mesolimbic innervation in two different mouse models either overexpressing or deficient for Reelin. In previous studies we reported that TgRln mice were more resilient to stressors implicated in the genesis of psychiatric diseases (chronic stress and psychostimulant administration) (Teixeira et al., 2011), suggesting a role for Reelin in preventing behavioral symptoms related with these disorders. Here we show that Reelin-depletion at adult stages does not lead to significant changes neither in the striatal composition nor in the dopaminergic innervation, while postnatal Reelin overexpression increases interneuron populations as well as the density of dopaminergic striatal projections from the VTA. Together, our results suggest the participation of postnatal Reelin expression in the fine structural tuning of the striatal area (Figure 7).

**Figure 7.**
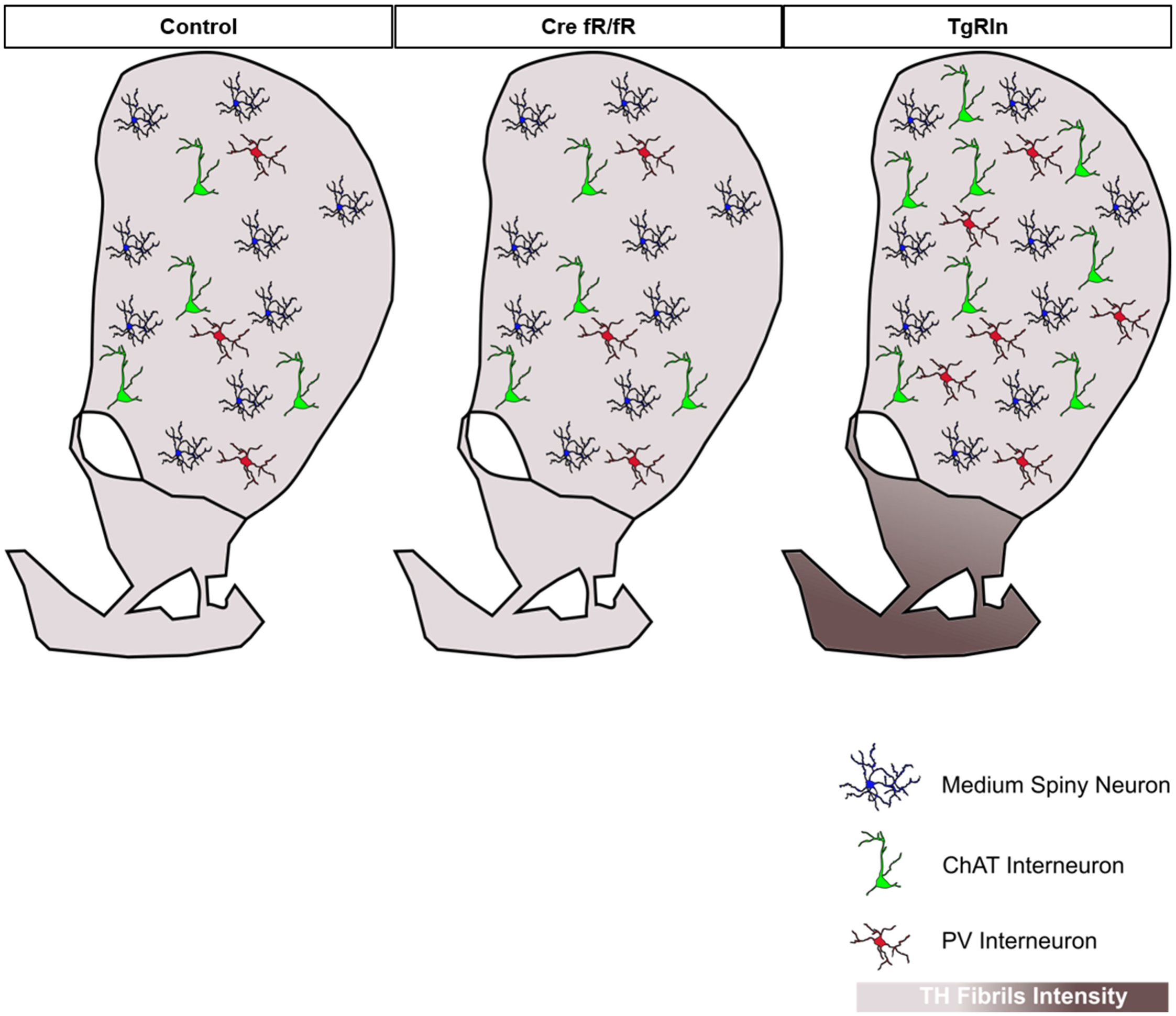
Schematic summary of the striatal organization in different Reelin mouse models. Density of striatal MSNs is preserved between the control, Cre fR/fR and TgRln striatums. Although the density of striatal PV-positive and ChAT-positive interneurons is maintained between control and Cre fR/fR mice, it is increased in the DS of TgRln mice. Increased numbers of ChAT-positive interneurons are present in the Dorsal striatum and higher numbers of PV-positive interneurons are distributed in the Ventral Medial striatum sub-region. Dopaminergic projections are represented with different gradient of brown colour, showing a specific increase of TH fibrils in the NAcc and OT of the TgRln mice compared with controls.

### 4.1 A role for Reelin in the striatum

The role of Reelin in the cortex and the hippocampus has been extensively studied including the expression pattern in GABAergic interneurons and the regulation in glutamatergic synapses (Alcántara et al., 1998; Herz and Chen, 2006; Jossin, 2020). Indeed, it has been characterized that Reelin controls several structural and functional properties of the glutamatergic synapses including the strength of glutamate neurotransmission (Beffert et al., 2005; Qiu et al., 2006b), protein composition at presynaptic boutons (Hellwig et al., 2011), structural properties of dendritic spines (Bosch et al., 2016) as well as trafficking of glutamate receptor subunits (Sinagra et al., 2005; Groc et al., 2007). Several studies also support a key role of Reelin in the correct organization of the basal ganglia. For instance, blockade of Reelin or its signaling pathway leads to a severe disorganization of the tangentially migrating midbrain dopaminergic (mDA) neurons, which fail to reach their final position in the SN pars compacta (SNc) and accumulate instead in the VTA. This results in a conspicuous reduction of the number of mDA neurons in the SNc, despite no overall changes in the number of mDA neurons have been described (Nishikawa et al., 2003; Kang et al., 2010; Sharaf et al., 2013; Bodea et al., 2014). Interestingly, alterations in the radial and tangential fibers that guide migrating mDA neurons have been described in *reeler* mice (Nishikawa et al., 2003; Kang et al., 2010) and support the idea that Reelin might also be guiding mDA neuronal migration indirectly by controlling the normal development of guidance scaffolds. However, specific inactivation of Reelin signaling in mDA neurons indicates a direct role of Reelin in the tangential migration of this neuronal population towards the SNc by promoting fast-laterally-directed migration and stabilization of their leading process (Vaswani et al., 2019). Despite these organization abnormalities in the SNc, no significant alterations have been described in the nigrostriatal pathway of *reeler, reeler-like* mutants or heterozygous reeler mice (Nishikawa et al., 2003; Sharaf et al., 2013; Vaswani et al., 2019). In contrast, defects in cortico-striatal plasticity (Marrone et al., 2006) and in the dopaminergic system (Matsuzaki et al., 2007) have been reported in *reeler* mice. Moreover, alterations in striatal composition, such as reductions in the number of striatal PV+ neurons along the rostro-caudal axis (Marrone et al., 2006; Ammassari-Teule et al., 2009), decreases in TH immunoreactivity in the NAcc (Nullmeier et al., 2014) and increases in the density of ChAT (Sigala et al., 2007) and the expression of D1, D2 and serotonin 5-HT2A receptors (Matsuzaki et al., 2007; Varela et al., 2015) when Reelin levels are decreased, have been also described.

In this study we describe a preferential expression of Reelin mRNA in a specific subpopulation of MSNs of the striatum, the D1 neurons, corroborating previous studies using FISH (de Guglielmo et al., 2022). Further, the fact that both Reelin and Dab1 expression are higher in striatonigral D1 MSNs than in striatopallidal D2 MSNs, suggests that Reelin may function in an autocrine manner in D1 MSNs.

Nevertheless, the lack of Reelin during development does not lead to dramatic alterations in the striatum of *reeler* mice. Considering the low Reelin expression levels in the midbrain and the profound defects associated with its absence, it has been hypothesized that Reelin may not act only by simple diffusion but also by axonal transport to target other brain structures (Nishikawa et al., 2003). The possibility that Reelin may be transported from a region such as the striatum to the SN or VTA is thus feasible and may represent the primary source of Reelin for midbrain neurons. Indeed, the idea that Reelin is anterogradely transported through striatonigral fibers of D1 MSNs to act on dopaminergic neurons seems to be relevant during migration but uncertain in adulthood since Reelin canonical receptors (i.e. ApoER2 and VLDLR) are not expressed in the adult midbrain (Sharaf et al., 2015). Considering the previous data, the specific effect of postnatal alterations of Reelin levels has been studied in detail in the striatum and interconnected areas including the SN and the VTA.

### 4.2 Consequences of the deficit of Reelin in psychiatric disorders

The description of heterozygous *reeler* mice as a useful model of psychosis vulnerability is still controversial since the phenotypic behavioral alterations observed could be attributable either to a role of Reelin during development or to an acute effect at adult stages. Given that very few studies have addressed this issue (Matsuzaki et al., 2007), here we use a conditional KO model (Cre fR/fR) in which neurodevelopment is preserved, which allowed us to specifically analyze the contribution of adult Reelin expression to the cellular and anatomical organization of the striatum. Previous studies have shown that in *reeler* and heterozygous-*reeler* mice there was a decrease in the density of PV+ cells in the Dorsal-Medial and Ventral-Medial striatal regions (Marrone et al., 2006; Ammassari-Teule et al., 2009). However, in Cre fR/fR mice we found no significant changes in cell densities of CHAT+ and PV+ interneurons. These differences can be attributable to the fact that in previous studies the lack of Reelin started during development, whereas in our study Reelin inactivation takes place at adult stages. In sum, these data suggest that Reelin expression is critical for striatal PV+ interneuron formation during striatal development, but not for maintenance of the pool of such interneuron populations during adulthood.

Similarly, previous studies evidenced alterations in TH expression in VTA and reduction in TH+ immunoreactivity terminals in striatum and VTA in heterozygous Reeler mice (Ballmaier et al., 2002). To analyze the effect of adult Reelin depletion, we mapped TH+ immunoreactivity in striatum, VTA and NAcc areas in Cre fR/fR mice, finding no differences with controls, although there was a trend in the OT. Our data suggests that at adult stages Reelin is largely dispensable for the maintenance of the dopaminergic innervation from the SN/VTA to the striatum.

### 4.3 Reelin overexpression in the striatum and drug sensitization

Although it has been widely described that the mesolimbic system controls drug sensitization, there are studies involving other striatal elements, such as the striatal patch-matrix organization and striatal interneurons, in the control of this process. We already reported that Reelin overexpression leads to reduced sensitization to cocaine (Teixeira et al., 2011). The characterization of the striatal organization in TgRln mice is essential to further understand the mechanisms underlying drug sensitization. Despite the fact that the gross structure of the striatal architecture was not altered in TgRln mice, the study of striatal interneurons, which represent 5% of the striatal cell population, clearly suggests that Reelin is able to modulate interneuron densities. For instance, Reelin overexpression leads to increased densities of PV+ and ChAT+ cells, suggesting a specific response of these neurons to increased amounts of Reelin. In addition, our results clearly indicate an increase of dopaminergic fibers in the NAcc and Olfactory tubercle of the TgRln mice. These alterations found in TgRln mice, but not in Cre fR/fR mice, reinforce the notion that increased Reelin levels modulate the striatal cytoarchitecture while Reelin presence in adulthood is not essential for the maintenance of the striatal organization.

Interestingly, decreased density of PV+ interneurons in the dorsomedial and ventromedial striatum of heterozygous *reeler* mice have been paralleled with deficits in some behaviors strongly disrupted in schizophrenic patients (Ammassari-Teule et al., 2009). Moreover, cocaine sensitization correlates with transient increases in the number of PV+ neurons in striatum that become reduced beyond normality after a 2-week cocaine withdrawal period (Todtenkopf et al., 2004). The fact that TgRln mice, which show reduced sensitization to cocaine, also show increased densities of PV+ interneurons could be apparently contradictory; nevertheless, here the number of PV+ interneurons is sustained, while upon cocaine administration the increase is transient, and eventually, related to compensatory responses. Anyhow, the fact that changes in PV+ interneuron number are controlling cocaine sensitization suggests that the increased density of PV+ cells observed in TgRln mice could be involved in the reduction of cocaine sensitization described in these mice (Teixeira et al., 2011). Although the mechanisms by which Reelin overexpression leads to increased numbers of PV and CHAT neurons remain unknown, it is important to remark that CAMKII promoter drives expression of Reelin in the striatum from the end of the first postnatal week onwards. It is thus possible that Reelin influences positively the maturation and survival of these interneurons, through Reelin/Dab1 associated pathways that influence these processes (Simó et al., 2007; Lee et al., 2014).

### 4.4 Molecular mechanisms of the effect of Reelin in the mesolimbic system

Disturbances in the dopaminergic mesolimbic system including altered immunoreactivity and mRNA levels of TH and dopamine transporters (D2, D3) in the VTA and the ventral striatum have been reported in heterozygous *reeler* mice (Ballmaier et al., 2002) and could be related to some of the behavioral deficits observed in this model. It has been described that after cocaine administration, there is an specific increase in the ERK pathway in striatonigral MSNs (Bertran-Gonzalez et al., 2008), a pathway that is also activated by Reelin (Simó et al., 2007; Lee et al., 2014). Interestingly, an increased Fos activation in the dorsal medial striatum but not in the NAcc of heterozygous reeler mice after the administration of cocaine has been described (de Guglielmo et al., 2022). Increases in Fos activation are thought to be the result of the cocaine-induced upregulation in dopamine levels in the striatum (Di Chiara and Imperato, 1988) which is hypothesized that might increase the activity of MSNs by activating D1 and D2 receptors. Experiments in mice lacking D1 receptor evidence a clear role for this receptor in the psychomotor effects of cocaine. As mentioned before, our data and that of others (de Guglielmo et al., 2022) evidences a preferential expression of Reelin in D1 neurons, supporting the idea that Reelin could be somehow modulating its function and hence influencing cocaine-induced psychomotor effects which are reduced in Reelin overexpressing mice (Teixeira et al., 2011) and increased when Reelin levels are reduced (de Guglielmo et al., 2022).

Specific Reelin activation in striatal neurons has not been proved so far, and additionally it has been described that expression of Reelin canonical receptors ApoER2 and to a lesser extent VLDLR is reduced in mature midbrain and striatum. From this data it can be assumed that Reelin functions are mostly restricted to migratory events and early postnatal maturation and that it is dispensable for the maintenance of dopaminergic neurons. Nevertheless, the putative contribution of the non-canonical Reelin pathway in ERK activation (Lee et al., 2014) maintains the potentiality of Reelin as a relevant factor. Together, we propose that Reelin overexpression in striatonigral MSNs could be controlling the ERK pathway and its feedback modulation to down regulate some responses to drug abuse.

Also interesting is the fact that the mesolimbic system is critical to induce drug sensitization (for example amphetamine (Perugini and Vezina, 1994)) which leads to a higher expression of c-fos positive cells in striatal patches rather than in the matrix compartment (Graybiel et al., 1990). Since Reelin has been described to be selectively expressed in striatal patches (Alcántara et al., 1998; Nishikawa et al., 1999) and Reelin controls immediate-early gene expression including Egr-1, Arc and c-fos amongst others (Simó et al., 2007; Stritt and Knöll, 2010) we could not discard that higher expression of Reelin in TgRln mice triggers reduced drug sensitization through the mediation of c-fos levels in striatal patches. Exploration of this eventuality will require specific research.

### 4.5 Reelin as a possible therapeutic target for psychiatric diseases

Reelin has been placed as a top candidate gene associated with several neuropsychiatric diseases. This link is supported by several studies showing that Reelin levels are reduced in patients with schizophrenia, bipolar disorder and autistic spectrum disorder (Impagnatiello et al., 1998; Fatemi et al., 2000, 2001, 2005; Persico et al., 2001; Torrey et al., 2005; Grayson et al., 2005; Ruzicka et al., 2007; Ovadia and Shifman, 2011; Wang et al., 2014; Baek et al., 2015; Lammert and Howell, 2016). Schizophrenia, which presents behavioral sensitization, can be compared with phenotypes related to drug sensitization where the protective effect of Reelin overexpression has been demonstrated (Teixeira et al., 2011). Moreover, in schizophrenic patients it has been described that densities of ChAT+ cell profiles were significantly reduced in the caudate nucleus, the ventral striatum and in the striatum as a whole in the schizophrenic group (Holt et al., 1999). Thus, Reelin overexpression may potentially counteract cholinergic interneuron alterations in schizophrenic patients.

It is interesting to notice that the striatal changes observed in TgRln mice are opposite to those found in patients with Tourette’s syndrome which present a clear decrease in the density of PV+ and ChAT+ interneurons in the dorsal striatum with no alterations in the density and number of MSNs (Kataoka et al., 2010). In TgRln mice, increased densities of PV+ and ChAT+ striatal interneurons with no overall alterations in the density of MSNs have been described. Importantly, the etiology of Tourette’s syndrome is heterogeneous and complex, with unclear mechanistic contributions, but with apparent dysfunction of interneurons functioning (Rapanelli et al., 2017), making it hard to find an effective treatment. Noteworthy, GWAS studies have identified RELN genetic variants in Tourette’s syndrome (Li et al., 2012) and together with our findings in the TgRln model suggest that Reelin could be an attractive therapeutic approach to reverse the symptoms of this disorder, although altered Reelin expression or signaling should be explored in patients affected by Tourette’s syndrome. The finding that adult-depletion of Reelin does not provoke significant alterations in striatum, but that Reelin overexpression induces changes in interneuron populations and dopaminergic innervations, positions Reelin fragments or pharmacological tools as top candidates for being used in future therapies.

## 6 Author Contributions

ES contributed to conception and design of the study. MP, SG, EM, LP, AE, NM, AV and YM performed the experiments. MP, SG, EM, CA and YM analyzed the data and performed statistical analysis. AN contributed new reagents/analytic tools, MP, LP, YM and ES wrote the first draft of the manuscript. EM, AP, JG, CA and MV wrote sections of the manuscript. YM and ES reviewed and edited the final version. All authors contributed to manuscript revision, read, and approved the submitted version.

## 8 Acknowledgments

We thank Daniela Rossi and Ashraf Muhaisen for help in the management of mouse colonies.

## Data Availability Statement

The original contributions presented in the study are included in the article/supplementary material, further inquiries can be directed to the corresponding author/s.

## References

Alcántara, S., Ruiz, M., D’Arcangelo, G., Ezan, F., de Lecea, L., Curran, T., et al. (1998). Regional and cellular patterns of reelin mRNA expression in the forebrain of the developing and adult mouse. J. Neurosci. 18, 7779–7799. doi: 10.1523/JNEUROSCI.18-19-07779.1998.

Ammassari-Teule, M., Sgobio, C., Biamonte, F., Marrone, C., Mercuri, N. B., and Keller, F. (2009). Reelin haploinsufficiency reduces the density of PV+ neurons in circumscribed regions of the striatum and selectively alters striatal-based behaviors. Psychopharmacology (Berl). 204, 511–521. doi: 10.1007/s00213-009-1483-x.

Arnaud, L., Ballif, B. A., and Cooper, J. A. (2003). Regulation of protein tyrosine kinase signaling by substrate degradation during brain development. Mol. Cell. Biol. 23, 9293–9302. doi: 10.1128/MCB.23.24.9293-9302.2003.

Baek, S. T., Copeland, B., Yun, E.-J., Kwon, S.-K., Guemez-Gamboa, A., Schaffer, A. E., et al. (2015). An AKT3-FOXG1-reelin network underlies defective migration in human focal malformations of cortical development. Nat. Med. 21, 1445–1454. doi: 10.1038/nm.3982.

Ballif, B. A., Arnaud, L., Arthur, W. T., Guris, D., Imamoto, A., and Cooper, J. A. (2004). Activation of a Dab1/CrkL/C3G/Rap1 pathway in Reelin-stimulated neurons. Curr. Biol. 14, 606–610. doi: 10.1016/j.cub.2004.03.038.

Ballmaier, M., Zoli, M., Leo, G., Agnati, L. F., and Spano, P. (2002). Preferential alterations in the mesolimbic dopamine pathway of heterozygous reeler mice: an emerging animal-based model of schizophrenia. Eur. J. Neurosci. 15, 1197–1205. doi: 10.1046/j.1460-9568.2002.01952.x.

Beffert, U., Morfini, G., Bock, H. H., Reyna, H., Brady, S. T., and Herz, J. (2002). Reelin-mediated signaling locally regulates protein kinase B/Akt and glycogen synthase kinase 3beta. J. Biol. Chem. 277, 49958–49964. doi: 10.1074/jbc.M209205200.

Beffert, U., Weeber, E. J., Durudas, A., Qiu, S., Masiulis, I., Sweatt, J. D., et al. (2005). Modulation of Synaptic Plasticity and Memory by Reelin Involves Differential Splicing of the Lipoprotein Receptor Apoer2. Neuron 47, 567–579. doi: 10.1016/j.neuron.2005.07.007.

Benhayon, D., Magdaleno, S., and Curran, T. (2003). Binding of purified Reelin to ApoER2 and VLDLR mediates tyrosine phosphorylation of Disabled-1. Brain Res. Mol. Brain Res. 112, 33–45.

Bertran-Gonzalez, J., Bosch, C., Maroteaux, M., Matamales, M., Hervé, D., Valjent, E., et al. (2008). Opposing Patterns of Signaling Activation in Dopamine D1 and D2 Receptor-Expressing Striatal Neurons in Response to Cocaine and Haloperidol. J. Neurosci. 28.

Bodea, G. O., Spille, J.-H., Abe, P., Andersson, A. S., Acker-Palmer, A., Stumm, R., et al. (2014). Reelin and CXCL12 regulate distinct migratory behaviors during the development of the dopaminergic system. Development 141, 661–673. doi: 10.1242/dev.099937.

Bolam, J. P. (1984). Synapses of identified neurons in the neostriatum. Ciba Found. Symp. 107, 30–47.

Bolam, J. P., Hanley, J. J., Booth, P. A., and Bevan, M. D. (2000). Synaptic organisation of the basal ganglia. J. Anat., 527–42.

Bosch, C., Masachs, N., Exposito-Alonso, D., Martínez, A., Teixeira, C. M., Fernaud, I., et al. (2016). Reelin Regulates the Maturation of Dendritic Spines, Synaptogenesis and Glial Ensheathment of Newborn Granule Cells. Cereb. Cortex 26, 4282–4298. doi: 10.1093/cercor/bhw216.

Chen, Y., Beffert, U., Ertunc, M., Tang, T.-S., Kavalali, E. T., Bezprozvanny, I., et al. (2005). Reelin Modulates NMDA Receptor Activity in Cortical Neurons. J. Neurosci. 25, 8209–8216. doi: 10.1523/JNEUROSCI.1951-05.2005.

Cooper, J. A. (2008). A mechanism for inside-out lamination in the neocortex. Trends Neurosci. 31, 113–119. doi: 10.1016/j.tins.2007.12.003.

Crittenden, J. R., and Graybiel, A. M. (2011). Basal Ganglia disorders associated with imbalances in the striatal striosome and matrix compartments. Front. Neuroanat. 5, 59. doi: 10.3389/fnana.2011.00059.

D’Arcangelo, G., G. Miao G., Chen, S.-C., Scares, H. D., Morgan, J. I., and Curran, T. (1995). A protein related to extracellular matrix proteins deleted in the mouse mutant reeler. Nature 374, 719–723. doi: 10.1038/374719a0.

D’Arcangelo, G., Homayouni, R., Keshvara, L., Rice, D. S., Sheldon, M., and Curran, T. (1999). Reelin is a ligand for lipoprotein receptors. Neuron 24, 471–9.

de Guglielmo, G., Iemolo, A., Nur, A., Turner, A., Montilla-Perez, P., Martinez, A., et al. (2022). Reelin deficiency exacerbates cocaine-induced hyperlocomotion by enhancing neuronal activity in the dorsomedial striatum. Genes. Brain. Behav. 21, e12828. doi: 10.1111/gbb.12828.

Di Chiara, G., and Imperato, A. (1988). Drugs abused by humans preferentially increase synaptic dopamine concentrations in the mesolimbic system of freely moving rats. Proc. Natl. Acad. Sci. U. S. A. 85, 5274–5278. doi: 10.1073/pnas.85.14.5274.

Fatemi, S. H., Earle, J. A., and McMenomy, T. (2000). Reduction in Reelin immunoreactivity in hippocampus of subjects with schizophrenia, bipolar disorder and major depression. Mol. Psychiatry 5, 571,654-663. doi: 10.1038/sj.mp.4000783.

Fatemi, S. H., Kroll, J. L., and Stary, J. M. (2001). Altered levels of Reelin and its isoforms in schizophrenia and mood disorders. Neuroreport 12, 3209–15.

Fatemi, S. H., Reutiman, T. J., and Folsom, T. D. (2009). Chronic psychotropic drug treatment causes differential expression of Reelin signaling system in frontal cortex of rats. Schizophr. Res. 111, 138–52. doi: 10.1016/j.schres.2009.03.002.

Fatemi, S. H., Snow, A. V, Stary, J. M., Araghi-Niknam, M., Reutiman, T. J., Lee, S., et al. (2005). Reelin signaling is impaired in autism. Biol. Psychiatry 57, 777–787. doi: 10.1016/j.biopsych.2004.12.018.

Folsom, T. D., and Fatemi, S. H. (2013). The involvement of Reelin in neurodevelopmental disorders. Neuropharmacology 68, 122–135. doi: 10.1016/j.neuropharm.2012.08.015.

Gerfen, C. R. (1992). The neostriatal mosaic: multiple levels of compartmental organization. Trends Neurosci. 15, 133–9.

Gernert, M., Hamann, M., Bennay, M., Löscher, W., and Richter, A. (2000). Deficit of striatal parvalbumin-reactive GABAergic interneurons and decreased basal ganglia output in a genetic rodent model of idiopathic paroxysmal dystonia. J. Neurosci. 20, 7052–8.

Gong, S., Doughty, M., Harbaugh, C. R., Cummins, A., Hatten, M. E., Heintz, N., et al. (2007). Targeting Cre recombinase to specific neuron populations with bacterial artificial chromosome constructs. J. Neurosci. Off. J. Soc. Neurosci. 27, 9817–9823. doi: 10.1523/JNEUROSCI.2707-07.2007.

González-Billault, C., Del Río, J. A., Ureña, J. M., Jiménez-Mateos, E. M., Barallobre, M. J., Pascual, M., et al. (2005). A role of MAP1B in Reelin-dependent neuronal migration. Cereb. Cortex 15, 1134–1145. doi: 10.1093/cercor/bhh213.

Graybiel, A. M., Moratalla, R., and Robertson, H. A. (1990). Amphetamine and cocaine induce drugspecific activation of the c-fos gene in striosome-matrix compartments and limbic subdivisions of the striatum. Proc. Natl. Acad. Sci. U. S. A. 87, 6912–6916. doi: 10.1073/pnas.87.17.6912.

Graybiel, A. M., and Ragsdale, C. W. (1978). Histochemically distinct compartments in the striatum of human, monkeys, and cat demonstrated by acetylthiocholinesterase staining. Proc. Natl. Acad. Sci. U. S. A. 75, 5723–6.

Grayson, D. R., Jia, X., Chen, Y., Sharma, R. P., Mitchell, C. P., Guidotti, A., et al. (2005). Reelin promoter hypermethylation in schizophrenia. Proc. Natl. Acad. Sci. U. S. A. 102, 9341 LP–9346. doi: 10.1073/pnas.0503736102.

Groc, L., Choquet, D., Stephenson, F. A., Verrier, D., Manzoni, O. J., and Chavis, P. (2007). NMDA Receptor Surface Trafficking and Synaptic Subunit Composition Are Developmentally Regulated by the Extracellular Matrix Protein Reelin. J. Neurosci. 27, 10165 LP–10175. doi: 10.1523/JNEUROSCI.1772-07.2007.

Heiman, M., Schaefer, A., Gong, S., Peterson, J. D., Day, M., Ramsey, K. E., et al. (2008). A Translational Profiling Approach for the Molecular Characterization of CNS Cell Types. Cell 135, 738–748. doi: 10.1016/j.cell.2008.10.028.

Hellwig, S., Hack, I., Kowalski, J., Brunne, B., Jarowyj, J., Unger, A., et al. (2011). Role for Reelin in neurotransmitter release. J. Neurosci. Off. J. Soc. Neurosci. 31, 2352–2360. doi: 10.1523/JNEUROSCI.3984-10.2011.

Herkenham, M., and Pert, C. B. (1981). Mosaic distribution of opiate receptors, parafascicular projections and acetylcholinesterase in rat striatum. Nature 291, 415–8.

Herz, J., and Chen, Y. (2006). Reelin, lipoprotein receptors and synaptic plasticity. Nat. Rev. Neurosci. 7, 850–859. doi: 10.1038/nrn2009.

Hiesberger, T., Trommsdorff, M., Howell, B. W., Goffinet, A., Mumby, M. C., Cooper, J. A., et al. (1999). Direct binding of Reelin to VLDL receptor and ApoE receptor 2 induces tyrosine phosphorylation of disabled-1 and modulates tau phosphorylation. Neuron 24, 481–489. doi: 10.1016/s0896-6273(00)80861-2.

Hirota, Y., and Nakajima, K. (2017). Control of Neuronal Migration and Aggregation by Reelin Signaling in the Developing Cerebral Cortex. Front. cell Dev. Biol. 5, 40. doi: 10.3389/fcell.2017.00040.

Holt, D. J., Herman, M. M., Hyde, T. M., Kleinman, J. E., Sinton, C. M., German, D. C., et al. (1999). Evidence for a deficit in cholinergic interneurons in the striatum in schizophrenia. Neuroscience 94, 21–31.

Howell, B. W., Hawkes, R., Soriano, P., and Cooper, J. A. (1997). Neuronal position in the developing brain is regulated by mouse disabled-1. Nature 389, 733–737. doi: 10.1038/39607.

Howell, B. W., Herrick, T. M., and Cooper, J. A. (1999). Reelin-induced tyrosine [corrected] phosphorylation of disabled 1 during neuronal positioning. Genes Dev. 13, 643–8.

Impagnatiello, F., Guidotti, A. R., Pesold, C., Dwivedi, Y., Caruncho, H., Pisu, M. G., et al. (1998). A decrease of reelin expression as a putative vulnerability factor in schizophrenia. Proc. Natl. Acad. Sci. U. S. A. 95, 15718–23.

Jossin, Y. (2020). Reelin Functions, Mechanisms of Action and Signaling Pathways During Brain Development and Maturation. Biomolecules 10. doi: 10.3390/biom10060964.

Kang, W.-Y., Kim, S.-S., Cho, S.-K., Kim, S., Suh-Kim, H., and Lee, Y.-D. (2010). Migratory defect of mesencephalic dopaminergic neurons in developing reeler mice. Anat. Cell Biol. 43, 241–251. doi: 10.5115/acb.2010.43.3.241.

Kataoka, Y., Kalanithi, P. S. A., Grantz, H., Schwartz, M. L., Saper, C., Leckman, J. F., et al. (2010). Decreased number of parvalbumin and cholinergic interneurons in the striatum of individuals with Tourette syndrome. J. Comp. Neurol. 518, 277–91. doi: 10.1002/cne.22206.

Kempermann, G. (2008). The neurogenic reserve hypothesis: what is adult hippocampal neurogenesis good for? Trends Neurosci. 31, 163–169. doi: 10.1016/j.tins.2008.01.002.

Kim, H. M., Qu, T., Kriho, V., Lacor, P., Smalheiser, N., Pappas, G. D., et al. (2002). Reelin function in neural stem cell biology. Proc. Natl. Acad. Sci. 99, 4020–4025. doi: 10.1073/pnas.062698299.

Krueger, D. D., Howell, J. L., Hebert, B. F., Olausson, P., Taylor, J. R., and Nairn, A. C. (2006). Assessment of cognitive function in the heterozygous reeler mouse. Psychopharmacology (Berl). 189, 95–104. doi: 10.1007/s00213-006-0530-0.

Lammert, D. B., and Howell, B. W. (2016). RELN Mutations in Autism Spectrum Disorder. Front. Cell. Neurosci. 10, 84. doi: 10.3389/fncel.2016.00084.

Lee, G. H., Chhangawala, Z., von Daake, S., Savas, J. N., Yates, J. R., Comoletti, D., et al. (2014). Reelin induces Erk1/2 signaling in cortical neurons through a non-canonical pathway. J. Biol. Chem. 289, 20307–17. doi: 10.1074/jbc.M114.576249.

Li, M. J., Wang, P., Liu, X., Lim, E. L., Wang, Z., Yeager, M., et al. (2012). GWASdb: a database for human genetic variants identified by genome-wide association studies. Nucleic Acids Res. 40, D1047–54. doi: 10.1093/nar/gkr1182.

Marrone, M. C., Marinelli, S., Biamonte, F., Keller, F., Sgobio, C. A., Ammassari-Teule, M., et al. (2006). Altered cortico-striatal synaptic plasticity and related behavioural impairments in reeler mice. Eur. J. Neurosci. 24, 2061–2070. doi: 10.1111/j.1460-9568.2006.05083.x.

Matsuzaki, H., Minabe, Y., Nakamura, K., Suzuki, K., Iwata, Y., Sekine, Y., et al. (2007). Disruption of reelin signaling attenuates methamphetamine-induced hyperlocomotion. Eur. J. Neurosci. 25, 3376–3384. doi: 10.1111/j.1460-9568.2007.05564.x.

McCutcheon, R. A., Brown, K., Nour, M. M., Smith, S. M., Veronese, M., Zelaya, F., et al. (2021). Dopaminergic organization of striatum is linked to cortical activity and brain expression of genes associated with psychiatric illness. Sci. Adv. 7. doi: 10.1126/sciadv.abg1512.

Molnár, Z., Clowry, G. J., Šestan, N., Alzu’bi, A., Bakken, T., Hevner, R. F., et al. (2019). New insights into the development of the human cerebral cortex. J. Anat. 235, 432–451. doi: 10.1111/joa.13055.

Nishikawa, S., Goto, S., Hamasaki, T., Ogawa, M., and Ushio, Y. (1999). Transient and compartmental expression of the reeler gene product reelin in the developing rat striatum. Brain Res. 850, 244–8.

Nishikawa, S., Goto, S., Yamada, K., Hamasaki, T., and Ushio, Y. (2003). Lack of Reelin causes malpositioning of nigral dopaminergic neurons: evidence from comparison of normal and Reln(rl) mutant mice. J. Comp. Neurol. 461, 166–73. doi: 10.1002/cne.10610.

Niu, S., Yabut, O., and D’Arcangelo, G. (2008). The Reelin signaling pathway promotes dendritic spine development in hippocampal neurons. J. Neurosci. 28, 10339–10348. doi: 10.1523/JNEUROSCI.1917-08.2008.

Nullmeier, S., Panther, P., Frotscher, M., Zhao, S., and Schwegler, H. (2014). Alterations in the hippocampal and striatal catecholaminergic fiber densities of heterozygous reeler mice. Neuroscience 275, 404–419. doi: 10.1016/j.neuroscience.2014.06.027.

Olson, L., Seiger, A., and Fuxe, K. (1972). Heterogeneity of striatal and limbic dopamine innervation: highly fluorescent islands in developing and adult rats. Brain Res. 44, 283–8.

Ovadia, G., and Shifman, S. (2011). The Genetic Variation of RELN Expression in Schizophrenia and Bipolar Disorder. PLoS One 6, e19955. doi: 10.1371/journal.pone.0019955.

Persico, A. M., D’Agruma, L., Maiorano, N., Totaro, A., Militerni, R., Bravaccio, C., et al. (2001). Reelin gene alleles and haplotypes as a factor predisposing to autistic disorder. Mol. Psychiatry 6, 150–159. doi: 10.1038/sj.mp.4000850.

Perugini, M., and Vezina, P. (1994). Amphetamine administered to the ventral tegmental area sensitizes rats to the locomotor effects of nucleus accumbens amphetamine. J. Pharmacol. Exp. Ther. 270.

Pujadas, L., Gruart, A., Bosch, C., Delgado, L., Teixeira, C. M., Rossi, D., et al. (2010). Reelin regulates postnatal neurogenesis and enhances spine hypertrophy and long-term potentiation. J. Neurosci. 30, 4636–4649. doi: 10.1523/JNEUROSCI.5284-09.2010.

Qiu, S., Korwek, K. M., Pratt-Davis, A. R., Peters, M., Bergman, M. Y., and Weeber, E. J. (2006a). Cognitive disruption and altered hippocampus synaptic function in Reelin haploinsufficient mice. Neurobiol. Learn. Mem. 85, 228–242. doi: 10.1016/j.nlm.2005.11.001.

Qiu, S., Zhao, L. F., Korwek, K. M., and Weeber, E. J. (2006b). Differential reelin-induced enhancement of NMDA and AMPA receptor activity in the adult hippocampus. J. Neurosci. 26, 12943–12955. doi: 10.1523/JNEUROSCI.2561-06.2006.

Rapanelli, M., Frick, L. R., and Pittenger, C. (2017). The Role of Interneurons in Autism and Tourette Syndrome. Trends Neurosci. 530, 481–484. doi: 10.1016/j.tins.2017.05.004.

Rice, D. S., and Curran, T. (2001). Role of the reelin signaling pathway in central nervous system development. Annu. Rev. Neurosci. 24, 1005–1039. doi: 10.1146/annurev.neuro.24.1.1005.

Ruzicka, W. B., Zhubi, A., Veldic, M., Grayson, D. R., Costa, E., and Guidotti, A. (2007). Selective epigenetic alteration of layer I GABAergic neurons isolated from prefrontal cortex of schizophrenia patients using laser-assisted microdissection. Mol. Psychiatry 12, 385–397. doi: 10.1038/sj.mp.4001954.

Schiffmann, S. N., Jacobs, O., and Vanderhaeghen, J. J. (1991). Striatal restricted adenosine A2 receptor (RDC8) is expressed by enkephalin but not by substance P neurons: an in situ hybridization histochemistry study. J. Neurochem. 57, 1062–7.

Sharaf, A., Bock, H. H., Spittau, B., Bouché, E., and Krieglstein, K. (2013). ApoER2 and VLDLr are required for mediating reelin signalling pathway for normal migration and positioning of mesencephalic dopaminergic neurons. PLoS One 8, e71091. doi: 10.1371/journal.pone.0071091.

Sharaf, A., Rahhal, B., Spittau, B., and Roussa, E. (2015). Localization of reelin signaling pathway components in murine midbrain and striatum. Cell Tissue Res. 359, 393–407. doi: 10.1007/s00441-014-2022-6.

Sigala, S., Zoli, M., Palazzolo, F., Faccoli, S., Zanardi, A., Mercuri, N. B., et al. (2007). Selective disarrangement of the rostral telencephalic cholinergic system in heterozygous reeler mice. Neuroscience 144, 834–844. doi: 10.1016/j.neuroscience.2006.10.013.

Simó, S., Jossin, Y., and Cooper, J. A. (2010). Cullin 5 regulates cortical layering by modulating the speed and duration of Dab1-dependent neuronal migration. J. Neurosci. Off. J. Soc. Neurosci. 30, 5668–5676. doi: 10.1523/JNEUROSCI.0035-10.2010.

Simó, S., Pujadas, L., Segura, M. F., La Torre, A., Del Río, J. A., Ureña, J. M., et al. (2007). Reelin induces the detachment of postnatal subventricular zone cells and the expression of the Egr-1 through Erk1/2 activation. Cereb. Cortex 17, 294–303. doi: 10.1093/cercor/bhj147.

Sinagra, M., Verrier, D., Frankova, D., Korwek, K. M., Blahos, J., Weeber, E. J., et al. (2005). Reelin, very-low-density lipoprotein receptor, and apolipoprotein E receptor 2 control somatic NMDA receptor composition during hippocampal maturation in vitro. J. Neurosci. Off. J. Soc. Neurosci. 25, 6127–6136. doi: 10.1523/JNEUROSCI.1757-05.2005.

Smith, Y., Bevan, M. D., Shink, E., and Bolam, J. P. (1998). Microcircuitry of the direct and indirect pathways of the basal ganglia. Neuroscience 86, 353–87.

Soriano, E., and Del Río, J. A. (2005). The cells of cajal-retzius: still a mystery one century after. Neuron 46, 389–394. doi: 10.1016/j.neuron.2005.04.019.

Strasser, V., Fasching, D., Hauser, C., Mayer, H., Bock, H. H., Hiesberger, T., et al. (2004). Receptor clustering is involved in Reelin signaling. Mol. Cell. Biol. 24, 1378–86.

Stritt, C., and Knöll, B. (2010). Serum response factor regulates hippocampal lamination and dendrite development and is connected with reelin signaling. Mol. Cell. Biol. 30, 1828–1837. doi: 10.1128/MCB.01434-09.

Teixeira, C. M., Kron, M. M., Masachs, N., Zhang, H., Lagace, D. C., Martinez, A., et al. (2012). Cell-autonomous inactivation of the reelin pathway impairs adult neurogenesis in the hippocampus. J. Neurosci. 32, 12051–12065. doi: 10.1523/JNEUROSCI.1857-12.2012.

Teixeira, C. M., Martín, E. D., Sahún, I., Masachs, N., Pujadas, L., Corvelo, A., et al. (2011). Overexpression of Reelin prevents the manifestation of behavioral phenotypes related to schizophrenia and bipolar disorder. Neuropsychopharmacology 36, 2395–2405. doi: 10.1038/npp.2011.153.

Todtenkopf, M., Stellar, J., Williams, E., and Zahm, D.. (2004). Differential distribution of parvalbumin immunoreactive neurons in the striatum of cocaine sensitized rats. Neuroscience 127, 35–42. doi: 10.1016/j.neuroscience.2004.04.054.

Torrey, E. F., Barci, B. M., Webster, M. J., Bartko, J. J., Meador-Woodruff, J. H., and Knable, M. B. (2005). Neurochemical markers for schizophrenia, bipolar disorder, and major depression in postmortem brains. Biol. Psychiatry 57, 252–260. doi: 10.1016/j.biopsych.2004.10.019.

Tueting, P., Costa, E., Dwivedi, Y., Guidotti, A., Impagnatiello, F., Manev, R., et al. (1999). The phenotypic characteristics of heterozygous reeler mouse. Neuroreport 10, 1329–34.

Varela, M. J., Lage, S., Caruncho, H. J., Cadavid, M. I., Loza, M. I., and Brea, J. (2015). Reelin influences the expression and function of dopamine D2 and serotonin 5-HT2A receptors: a comparative study. Neuroscience 290, 165–174. doi: 10.1016/j.neuroscience.2015.01.031.

Vaswani, A. R., Weykopf, B., Hagemann, C., Fried, H.-U., Brüstle, O., and Blaess, S. (2019). Correct setup of the substantia nigra requires Reelin-mediated fast, laterally-directed migration of dopaminergic neurons. Elife 8. doi: 10.7554/eLife.41623.

Vílchez-Acosta, A., Manso, Y., Cárdenas, A., Elias-Tersa, A., Martínez-Losa, M., Pascual, M., et al. (2022). Specific contribution of Reelin expressed by Cajal-Retzius cells or GABAergic interneurons to cortical lamination. Proc. Natl. Acad. Sci. U. S. A. 119, e2120079119. doi: 10.1073/pnas.2120079119.

Wang, Z., Hong, Y., Zou, L., Zhong, R., Zhu, B., Shen, N., et al. (2014). Reelin gene variants and risk of autism spectrum disorders: An integrated meta-analysis. Am. J. Med. Genet. Part B Neuropsychiatr. Genet. 165, 192–200. doi: 10.1002/ajmg.b.32222.

Yasui, N., Nogi, T., and Takagi, J. (2010). Structural Basis for Specific Recognition of Reelin by Its Receptors. Structure 18, 320–331. doi: 10.1016/j.str.2010.01.010.

Zhao, C., Deng, W., and Gage, F. H. (2008). Mechanisms and Functional Implications of Adult Neurogenesis. Cell 132, 645–660. doi: 10.1016/j.cell.2008.01.033.

